# The *Mycobacterium tuberculosis* sRNA F6 modifies expression of essential chaperonins, GroEL2 and GroES

**DOI:** 10.1101/2020.07.15.204107

**Authors:** Joanna Houghton, Angela Rodgers, Graham Rose, Alexandre D’Halluin, Terry Kipkorir, Declan Barker, Simon J Waddell, Kristine B. Arnvig

**Affiliations:** Faculty of Infectious Tropical Diseases, London School of Hygiene and Tropical medicine, London WC1E 7HT, UK; Mycobacterial Metabolism and Antibiotic Research and Host-Pathogen Interactions in Tuberculosis Laboratories, The Francis Crick Institute, London NW1 1AT, UK; North Thames Genomic Laboratory Hub, Great Ormond Street Hospital for Children, London WC1N 3BH, UK; Global Health and Infection, Brighton and Sussex Medical School, University of Sussex, Brighton, BN1 9PX, UK; Structural and Molecular Biology, University College London, London WC1E 6BT, UK

**Keywords:** *Mycobacterium tuberculosis*, small RNA, nutrient starvation, Wayne model chaperonins, infection

## Abstract

Almost 140 years after the identification of *Mycobacterium tuberculosis* as the etiological agent of tuberculosis, important aspects of its biology remain poorly described. Little is known about the role of post-transcriptional control of gene expression and RNA biology, including the role of most of the small RNAs (sRNAs) identified to date. We have carried out a detailed investigation of the *M. tuberculosis* sRNA, F6, and shown it to be dependent on SigF for expression, and significantly induced in starvation conditions *in vitro* and in a mouse model of infection. Further exploration of F6 using an *in vitro* starvation model of infection indicates that F6 affects the expression of the essential chaperonins, GroEL2 and GroES. Our results point towards a role for F6 during periods of low metabolic activity typically associated with long-term survival of *M. tuberculosis* in human granulomas.

## INTRODUCTION

One hundred years after the launch of the Bacillus Calmette-Guerin (BCG) vaccine against tuberculosis (TB), this disease still claims more than 3000 lives on a daily basis, and its aetiological agent, *Mycobacterium tuberculosis* remains one of the most prominent pathogens in human history. Notoriously difficult to work with and very different from both Gram-positive and Gram-negative model organisms, many aspects of its basic biology, including RNA biology and post-transcriptional control of gene expression remain unclear. However, the study of regulatory RNA in *M. tuberculosis* is gaining momentum, aided by next-generation sequencing (NGS) applications, which have provided significant insights into the abundance and dynamics of non-coding RNA over a range of growth conditions, the location of transcription start sites on a global scale and translated *versus* untranslated transcripts (1–5). Still, much remains to be uncovered about the post-transcriptional control exerted by regulatory RNA in *M. tuberculosis* in particular what role these molecules play in the pathogenesis and persistence of *M. tuberculosis*. A multitude of *M. tuberculosis* small regulatory RNAs (sRNAs) have been identified in the last decade, but only few have been investigated and even fewer, linked to *Mtb* persistence and TB latency (1, 3, 4, 6–13).

F6 (ncRv10243) was one of the first *M. tuberculosis* sRNAs to be identified by cDNA cloning rather than RNA-seq (6). F6 is conserved in a wide range of mycobacteria with the 5’ end showing the highest degree of conservation (6, 7). The location of F6 between the convergent *fadA2* (encoding an acetyl-CoA transferase) and *fadE5* (encoding an acyl-CoA hydrogenase), is also highly conserved suggesting a potential role for F6 in the regulation of lipid metabolism, which is critical for intracellular survival (14). Initial analysis, combining northern blotting with 5’ and 3’ RACE, revealed that F6 is expressed as a 102-nucleotide transcript, which is 3’ processed to the dominant transcript of 58 nucleotides (6). F6 is upregulated in stationary phase, during oxidative stress, and by low pH, while overexpression from a multicopy-number plasmid leads to a slow-growth phenotype on solid media (6). The F6 promoter contains a typical SigF promoter motif and exhibits the highest SigF occupancy according to ChIP-chip analysis (6, 15). SigF is a non-essential sigma factor, conserved in most mycobacteria; its expression is induced by several stresses including anaerobiosis, nutrient starvation, oxidative stress and cold shock, while heat shock downregulates its expression, and deletion leads to attenuation in mice (16–19).

In the current study, we have investigated *M. tuberculosis* F6 expression under different growth conditions and found that this sRNA is significantly upregulated in a nutrient starvation model of persistence and in mouse lungs. We have generated an F6 deletion strain of *M. tuberculosis* and assessed the fitness of the Δ*f6 M. tuberculosis* strain *in vitro* and *in vivo*. We found that inactivation of F6 leads to impaired recovery from a Wayne model of hypoxia. Moreover, using microarray analysis and qRT-PCR, we have found that expression of the essential chaperonins *groEL2/rv0440* and *groES/rv3418c* were significantly upregulated upon deletion of F6. Our results suggest that although F6 is highly upregulated in *M. tuberculosis* during the early stages of infection, it is likely to play a more prominent role during later stages of infection.

## MATERIALS AND METHODS

### Bacterial strains and plasmids

*Escherichia coli* DH5α was used for plasmid construction and grown in LB agar or broth using kanamycin at 50 μg/ml and X-Gal where necessary at 200 μg/ml. *Mycobacterium tuberculosis* H37Rv was grown on Middlebrook 7H11 agar plus 10% OADC (Becton Dickinson). Liquid cultures were grown in standard Middlebrook 7H9 medium supplemented with 0.5% glycerol, 10% Middlebrook ADC (Becton Dickinson) and 0.05% Tween-80 at 37°C in a roller bottle (Nalgene) rolling at 2 rpm or 50 ml falcon tubes (Corning) in an SB3 Tube Rotator (Stewart) at 28 rpm. Kanamycin was added where required at 25 μg/ml and X-Gal where required at 50 μg/ml. All plasmids and oligos used in this study are listed in Tables S1 and S2.

### The Wayne Model

Cultures were grown to exponential phase (OD_600_=0.6-0.8) and subsequently diluted to OD_600_=0.005 in 7H9 in triplicate in Wayne tubes, each containing a sterile stirring bar. Cultures were incubated at 37°C on a stirring platform and OD monitored over time until cultures reached NRP-2. Cultures were then either diluted into 7H9 OD_600_=0.05 and OD monitored, or serially diluted onto 7H11 agar and CFU enumerated.

### Construction of F6 deletion and complemented strains

Allelic replacement techniques were used to generate an *M. tuberculosis* knockout mutant as per published protocol (20). Briefly, approximately 1.5 kb of the flanking region from either side of F6 (5’ flank coordinates 292520-293605 and 3’ flank 293709-294600) were cloned into the suicide vector pBackbone (21). The sRNA was replaced by an Xba I site using site directed mutagenesis to produce the targeting plasmid pJHP04. Electrocompetent H37Rv was transformed with pJHP04 and subsequent single cross overs (SCOs) and double cross overs (DCOs) selected (see supplementary material).

To complement the Δ*f6* mutant, electrocompetent cells were prepared and transformed with an integrating plasmid (pJHP06) containing the cloned region 293428-293876 from *M. tuberculosis* H37Rv, supplemented with pBSInt, providing the phage integrase (22). Complementation was confirmed using qRT-PCR.

### Mapping genome data and variant calling

To determine the genome sequence of the deletion strain, WGS was performed as described elsewhere using the Illumina HiSeq platform (2). Sequencing reads for the Δ*f6* strain were mapped against the *M. tuberculosis* H37Rv reference genome (AL123456) using BWA (23), and variants called with SAMtools (24). Variant filtering was performed by inclusion of only those variants with a minimum mapping quality of 10 and maximum read depth of 400. Finally, heterozygous calls or those found in repetitive or mobile elements (genes annotated as PE/PPE/insertions/phages) were removed.

### Preparation of starved cultures

*M. tuberculosis* H37Rv, Δ*f6* and complemented strain were grown in Middlebrook 7H9 supplemented with 0.4% glycerol, 0.085% NaCl, 0.5% BSA and 0.05% Tyloxapol in roller bottle culture (2 rpm at 37°C). Exponentially growing bacteria were pelleted and washed 3 times with PBS supplemented with 0.025% Tyloxapol and finally resuspended in triplicate in PBS with 0.025% Tyloxapol to an equivalent starting volume and incubated statically for 24 or 96 hours.

### RNA isolation

RNA isolation from *in vitro* cultures was done as described previously (6). Briefly, cultures were harvested with rapid cooling by adding ice directly to the culture and subsequent centrifugation at 10,000 rpm for 10 minutes. RNA was isolated from the pellet using the FastRNA Pro Blue Kit from MP Biomedicals following manufacturer’s instructions.

To isolate RNA from bacteria grown in mice, lung homogenates were spun at 13,000 rpm for 5 minutes to collect the bacteria. These were resuspended in 1 ml Trizol (Invitrogen) with 150-micron glass beads and the samples disrupted in a fast Prep (MP Biomedicals) at a setting of 6.0 for 40 secs. The RNA was extracted according to manufacturer’s guidelines. RNA concentration was measured by Nanodrop (Thermo Scientific) and RNA integrity measured by the 2100 Bioanalyzer using a Nano chip (Agilent Technologies).

### Quantitative RT-PCR

Total RNA was treated with Turbo DNase (Ambion) until DNA free. cDNA was synthesized using Superscript III (Invitrogen) and random hexamers. Primers were designed using the Applied Biosystems software Primer Express, and sequences are listed in Table S1. Each 20 μl qRT-PCR reaction, contained 16SYBRgreen (Applied Biosystems), 900 nm each primer and 5 μl of template cDNA. Absolute quantitation was carried out and all genes were normalised to 16S rRNA expression.

### Transcriptional profiling

Whole genome *M. tuberculosis* microarray slides were purchased from Agilent Technologies through the Bacterial Microarray Group at St. George’s (BμG@S), University of London. For cDNA synthesis, 2 μg wildtype H37Rv and Δ*sfdS* knockout, isolated from 24 hour starved cultures was used. The cDNA was labelled individually with both Cy-3 and Cy-5 dyes (GE Healthcare) using Superscript III reverse transcriptase (Invitrogen). Dye swaps were performed, and the cDNA hybridised to an 8 Chamber Agilent slide at 65°C for 16 hours before washing the slide with Oligo aCGH Wash Buffer 1 (Agilent) for 5 minutes at room temperature and Oligo aCGH Wash Buffer 2 (Agilent) for 1 minute at 37°C. Slides were stabilised using Agilent Stabilisation and Drying Solution according to manufacturer’s instructions.

Slides were scanned at 5 microns using an Agilent Technologies Microarray Scanner at BμG@S. Txt files created by the Agilent scanner were analysed using Genespring 14.5 filtering on flags and expression. T-test against zero was performed using p-value p<0.05with Benjamini-Hochberg multiple testing correction and 2-fold cut-off. Array design is available in BμG@Sbase (Accession No. A-BUGS-41; http://bugs.sgul.ac.uk/A-BUGS-41) and ArrayExpress (Accession No. A-BUGS-41). Microarray data have been deposited into ArrayExpress (Accession number E-MTAB-9327).

### F6 Target prediction

Prediction of F6 sRNA binding targets was performed using the TargetRNA2 webserver (http://cs.wellesley.edu/~btjaden/TargetRNA2/index.html) (25). Targets were predicted from the entire H37Rv genome sequence using full-length F6 and default parameters.

### Macrophage Infection

Bone marrow derived macrophages (BMDMs) were generated from 6-8 week old BALB/c mice in RPMI-1640 (Gibco) containing 10% foetal calf serum, 20 μM L-glutamine, 1 mM sodium pyruvate, 10 μM HEPES and 50 nM β-mercaptoethanol. The cells were then grown and differentiated in complete RPMI-1640 supplemented with 20% L929 cell supernatant for 6 days at 37°C in 5% CO_2_. The differentiated cells were seeded at a density of 2 x 10^5^ cells/well in 1 ml complete RPMI-1640 supplemented with 5% L929 cell supernatant and incubated overnight prior to infection. *M. tuberculosis* strains for infection were grown to an OD_600_ of 0.5-0.8 and inocula prepared by washing and resuspending the cultures in PBS, to produce a single cell suspension. This was used to infect BMDMs at a multiplicity of infection (MOI) of 0.1:1. After 4 hr, the cells were washed to remove all extracellular bacilli, medium replaced and incubation continued. Macrophages were lysed with water-0.05% Tween 80 to release intracellular bacteria after 4, 24, 72, 120, 168 hr post-infection. Bacilli were serially diluted in PBS-Tween and plated on 7H11 with OADC. Plates were incubated for 3-4 weeks for CFU counts. All experiments were performed in triplicate.

### Murine infection model and ethics statement

Groups of 6–8 week old BALB/c mice were infected by low-dose aerosol exposure with *M. tuberculosis* H37Rv wildtype, Δ*f6* and the complemented strain using a Glas-Col (Terre Haute, IN) aerosol generator calibrated to deliver approximately 100 bacteria into the lungs. Bacterial counts in the lungs (n = 5) at each time point of the study were determined by plating serial dilutions of individual lung homogenates on duplicate plates of Middlebrook 7H11 agar containing OADC enrichment. Colony-forming units were counted after 3–4 weeks incubation at 37°C. BALB/c mice were bred and housed under specific pathogen-free conditions at the Medical Research Council, National Institute for Medical Research (NIMR). All mouse studies and breeding were approved by the animal ethics committee at NIMR. Protocols for experiments were performed, under project license number 80/2236, in accordance with Home Office (United Kingdom) requirements and the Animal Scientific Procedures Act, 1986.

## RESULTS

### Expression of F6/SfdS is upregulated by starvation and during infection

SigF is the predicted main regulator of F6 expression, and this was validated by northern blotting, which demonstrated a complete absence of F6 in a Δ*sigF* strain (Fig. S1). We will henceforth refer to F6 as <u>S</u>fdS for Sig<u>F</u> <u>d</u>ependent <u>s</u>RNA. SigF is highly expressed during infection and in persister *in vitro* models such as nutrient starvation obtained by static incubation in PBS (16). We subjected *M. tuberculosis* H37Rv to starvation by washing and resuspending log-phase cultures in PBS for 24 and 96 hours before isolating total RNA. SfdS expression was measured by qRT-PCR (normalised to 16S rRNA) and compared to log-phase levels.

The results demonstrate that SfdS expression increased dramatically (19-fold) within 24 hours of starvation, with levels of SfdS remaining elevated for at least another 72 hours (Fig. 1).

**Figure 1:**
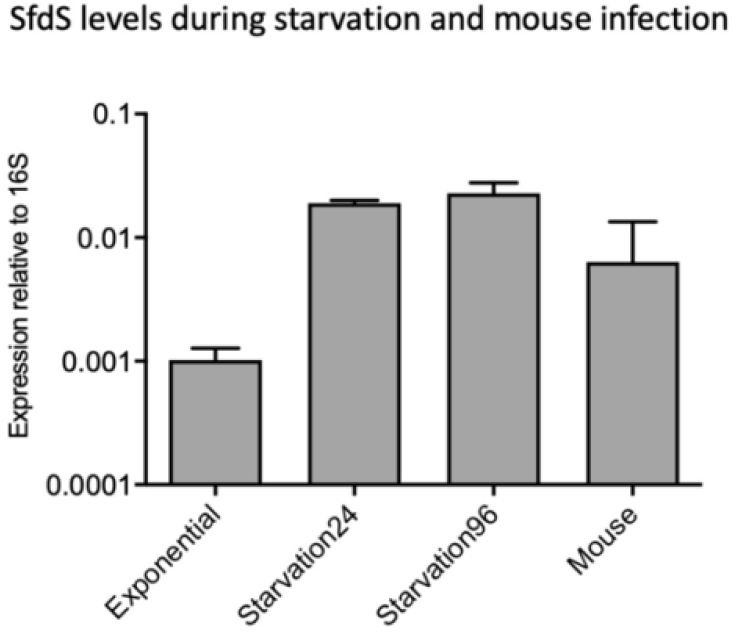
Expression of SfdS in *M. tuberculosis*. SfdS expression levels were measured in exponentially growing cultures, after 24 and 96 hours in PBS (Starvation24 and Starvation96) and in mouse lungs after three week’s infection using quantitative RT-PCT (qRT-PCR). Expression levels were normalised to 16S rRNA and the data represents the mean and standard deviation of three biological replicates for each condition.

To investigate SfdS expression in a mouse model of infection, BALB/c mice were infected with ~100 CFU of H37Rv via aerosol route and left for three weeks before culling and isolation of *M. tuberculosis* total RNA from lung tissue. We found that SfdS expression was robustly up-regulated (~ 6-fold) compared to log-phase levels, but not to the same extent as during *in vitro* starvation (Fig. 1).

Together these results demonstrate that the expression of SfdS is dynamic and that it may play a role during infection, more specifically in nutrient-poor environments.

### Deletion of SfdS does not impair *M. tuberculosis* standard growth *in vitro*

To identify a potential regulatory role of SfdS with minimal impact on the two flanking *fad* genes, we generated an unmarked SfdS deletion strain using site directed mutagenesis and allelic exchange. Candidates were screened by PCR amplification and sequencing of the PCR product, which confirmed the deletion of the sRNA from the *M. tuberculosis* genome (not shown).

To further verify the deletion and to determine if there were secondary mutations that might affect subsequent phenotypic analysis, wildtype and Δ*sfdS* strains were sequenced. Alignment of the genomic sequence of Δ*sfdS* with H37Rv identified two single-nucleotide polymorphisms (SNPs) in addition to the Xba I site deliberately introduced; one was C2864730T, resulting in an R102W substitution in Rv2541 (hypothetical protein) while the other was C4178146T in the 5’ UTR of Rv3729 (potential transferase). To ensure that these SNPs did not influence the phenotype of the SfdS deletion strain, we constructed a complemented strain, in which expression of SfdS was driven by its native promoter from a single-copy plasmid integrated on the chromosome. Growth of wildtype H37Rv, Δ*sfdS*, and the complemented strain was monitored over a period of two weeks in 7H9 roller bottle cultures. Under these conditions, there was no significant difference between the three strains (Fig. 2).

**Figure 2:**
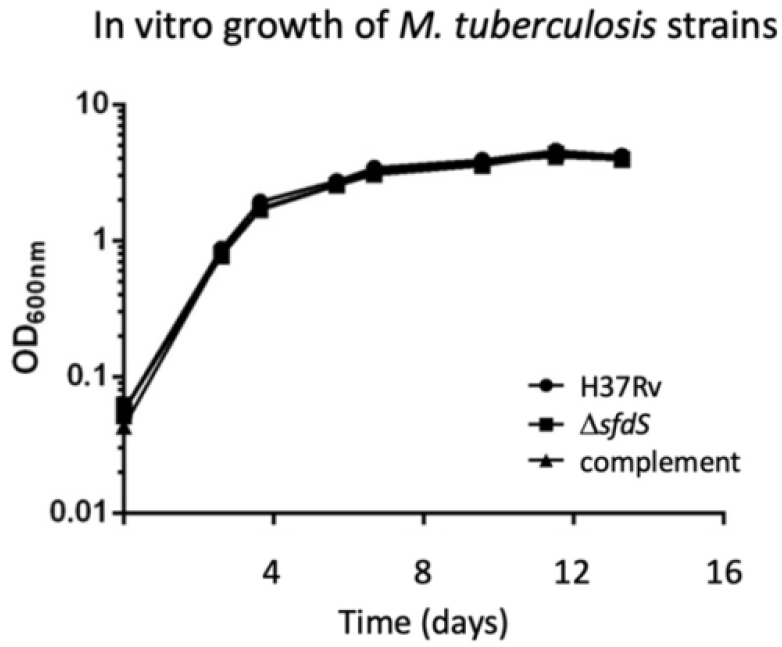
*In vitro* growth of Δ*sfdS*. Growth curves of wildtype *M. tuberculosis* H37Rv, Δ*sfdS* and its complement in 7H9 in roller bottles. The results represent the average and standard deviation of three biological replicates.

To validate deletion and complementation, expression of SfdS was measured in wildtype H37Rv, Δ*sfdS* and the complemented strain during exponential growth and after 24 hours of PBS starvation, using qRT-PCR. The results demonstrated that there was no detectable expression of SfdS in the deletion strain (Fig. S2), and that there was a small, non-significant difference between SfdS expression in wildtype and complemented strains in both growth conditions (unpaired t-test, p<0.05, Figure 3).

**Figure 3:**
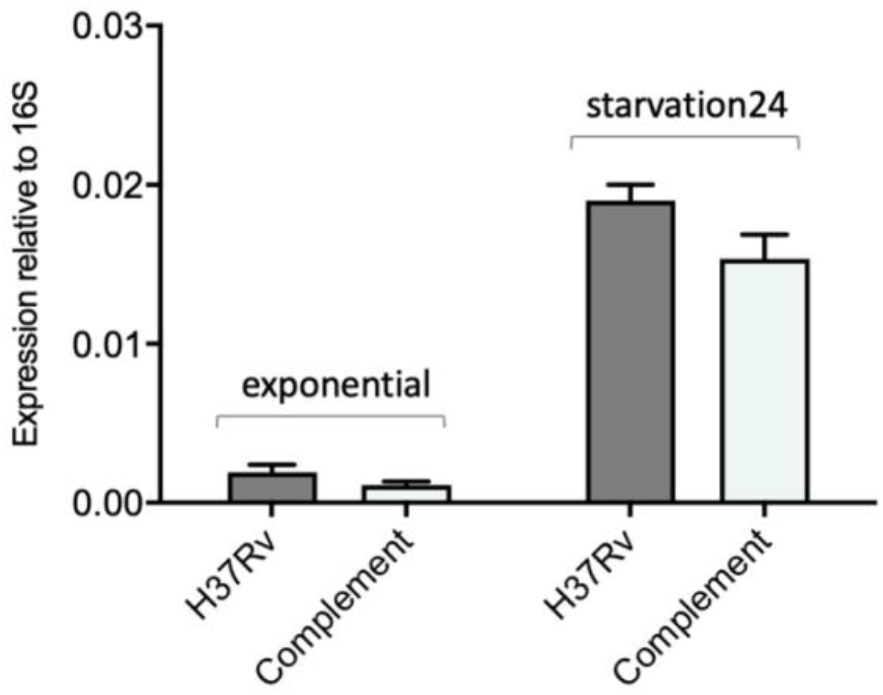
Expression of SfdS in wildtype H37Rv and complemented strain. RNA was isolated from log-phase and after 24 hours starvation and analysed by qRT-PCR. Data represents the mean and standard deviation of three biological replicates for each strain. Differences between wildtype and complement not significant according to unpaired t-test.

### Fitness of the Δ*sfdS* strain in macrophages and mouse models of infection

As we had observed a robust upregulation of SfdS during infection, we wanted to compare the fitness of wildtype *M. tuberculosis* H37Rv, Δ*sfdS* and the complemented strain during infection. The three strains were used to infect naïve and IFN-γ pre-activated murine BMDMs at an MOI of 0.1:1 (bacteria:macrophages). The infection was allowed to continue for 7 days with five time points. At each time point, the macrophages were lysed and plated for CFUs. For both naïve and activated macrophages, we observed no significant differences between the three strains, suggesting that under these conditions, deletion of SfdS does not result in attenuation (Fig. 4A).

**Figure 4:**
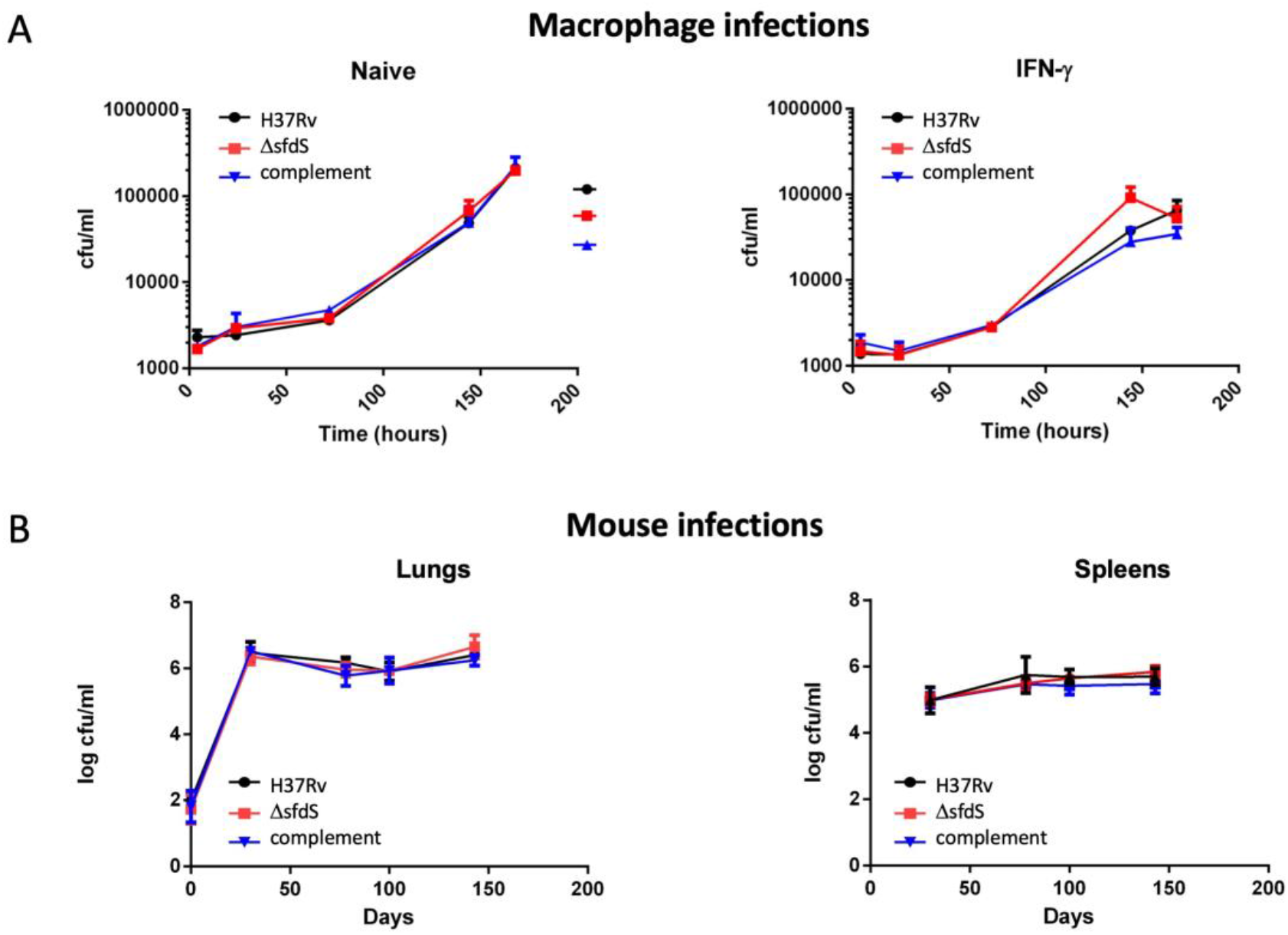
Survival of the Δ*sfdS* in models of infection. Panel A shows survival of wildtype *M. tuberculosis* H37Rv, ΔsfdS, and complementing strains in a macrophage model of infection with naive (left) and IFN-γ activated (right) BMDM. Data represent the mean and standard deviation of triplicate infections. Statistical significance was tested with One-way ANOVA p<0.05. Panel B shows survival of the three strains within the lungs and spleens of BALB/c mice. Data represents the averages and standard deviations from 5 mice per time point. Statistical significance was tested with Two-way ANOVA.

A macrophage model of infection does not entirely replicate the conditions encountered by the bacteria in more complex animal models of infection. To assess the role of SfdS in pathogenesis in a more representative infection model, wildtype *M. tuberculosis* H37Rv, Δ*sfdS*, and the complemented strain were compared in a mouse model of infection. BALB/c mice were infected with approximately 100 CFU via the aerosol route and the infection followed for 143 days. On days 0, 30, 78, 100 and 143, lungs and spleens were harvested for bacterial enumeration. Again, no significant difference was observed in CFUs between the wildtype *M. tuberculosis* H37Rv and Δ*sfdS* at any of the time points (Fig. 4B). The absence of an obvious phenotype is possibly due to the choice of mouse model and/or low MOI used in this experiment.

### Recovery from Wayne Model hypoxia is impaired in Δ*sfdS*

One of the hallmarks of human TB is the formation of granulomas, which are absent in a standard mouse model. Granulomas are characterised by limited nutrient and oxygen availability among other things (26). A widely used *in vitro*-model of hypoxia is the Wayne model, in which the available oxygen is gradually limited in sealed cultures of *M. tuberculosis* (27). When oxygen concentrations decrease to the microaerobic level (1% oxygen saturation), the cells enter a state of non-replicating persistence, NRP-1, followed by NRP-2 when oxygen saturation reaches 0.06%. SigF is highly induced during anaerobiosis, suggesting that its regulon, including SfdS, may play a role in low-oxygen conditions (19). We therefore decided to evaluate the fitness of the Δ*sfdS* strain using the Wayne model. To measure potential differences in respiration rate, the depletion of oxygen for each strain was monitored by a methylene blue indicator tube set up for each strain at the start of incubation. After 14 days incubation, at NRP-2, cultures were plated onto 7H11 agar for determination of colony forming units (CFU). In addition, NRP-2 cells were assessed for their ability to be resuscitated by dilution into fresh 7H9 media and monitoring of growth (by OD_600_). There was no significant difference in survival (CFU) between wildtype H37Rv, Δ*sfdS*, and the complemented *M. tuberculosis* strain at day 14 incubation in the Wayne model (Fig. 5A). Conversely, the re-growth of NRP-2 cells transferred to fresh media indicated that the Δ*sfdS* strain was impaired for recovery/resuscitation following incubation in the Wayne model, while the complemented strain displayed an intermediate phenotype (Fig. 5B). These results suggest that SfdS may play a role in the resuscitation of non-replicating *M. tuberculosis*, which is associated with the reactivation of latent infection.

**Figure 5:**
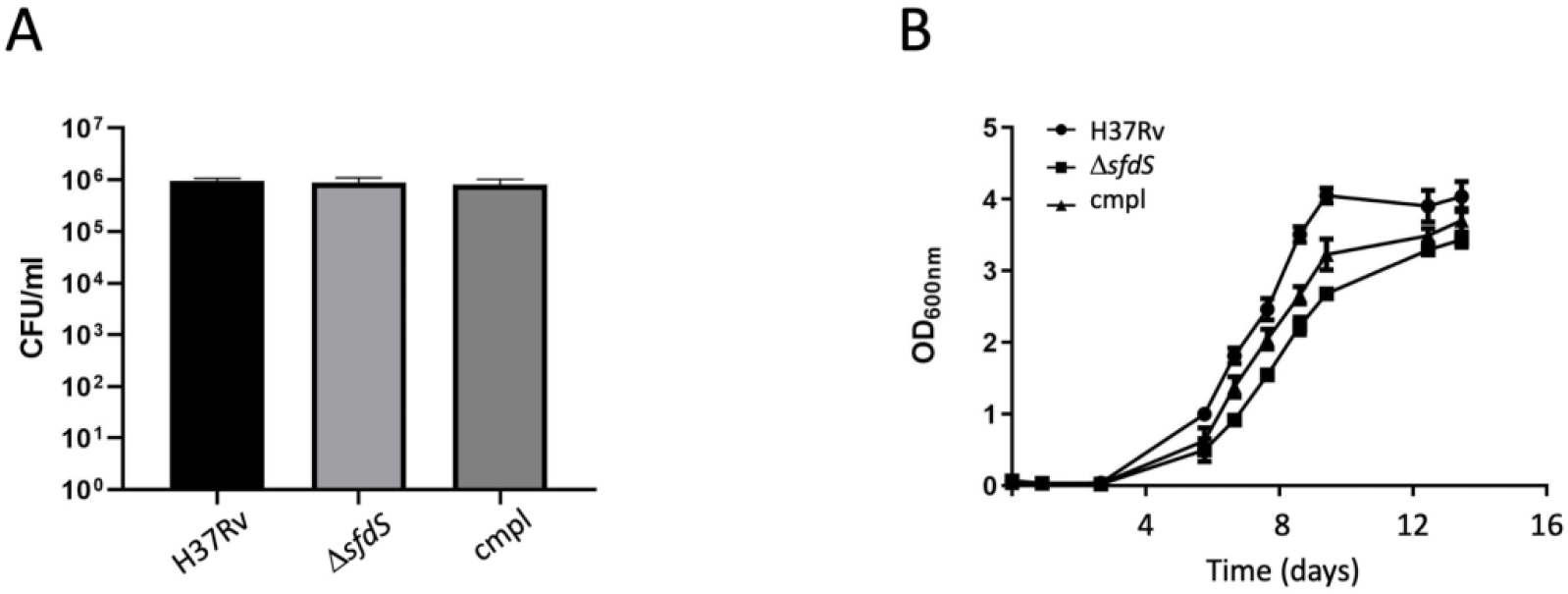
Recovery from NRP-2 is impaired in Δ*sfdS*. Wildtype *M. tuberculosis* H37Rv, Δ*sfdS* and complement (cmpl) strains were grown in the Wayne model until NRP-2 was reached. Cultures were diluted and plated for CFU counts. (A) Number of viable bacteria after incubation in the Wayne model. All cultures were subsequently adjusted to OD_600_=0.05 in fresh media and growth measured over 14 days (B). All data represents the averages and standard deviation of three biological replicates. Linear regression analysis indicated a significant difference (p<0.05) in the recovery of wildtype and Δ*sfdS*.

### Deletion of SfdS leads to upregulation of essential chaperonins

To establish a potential regulatory role for SfdS, we employed nutrient starvation, as this was the most potent inducer of the sRNA in our hands. RNA was isolated from triplicate cultures of wildtype H37Rv and Δ*sfdS* starved for 24 hours in PBS and analysed using Agilent microarrays.

**Figure 5:**
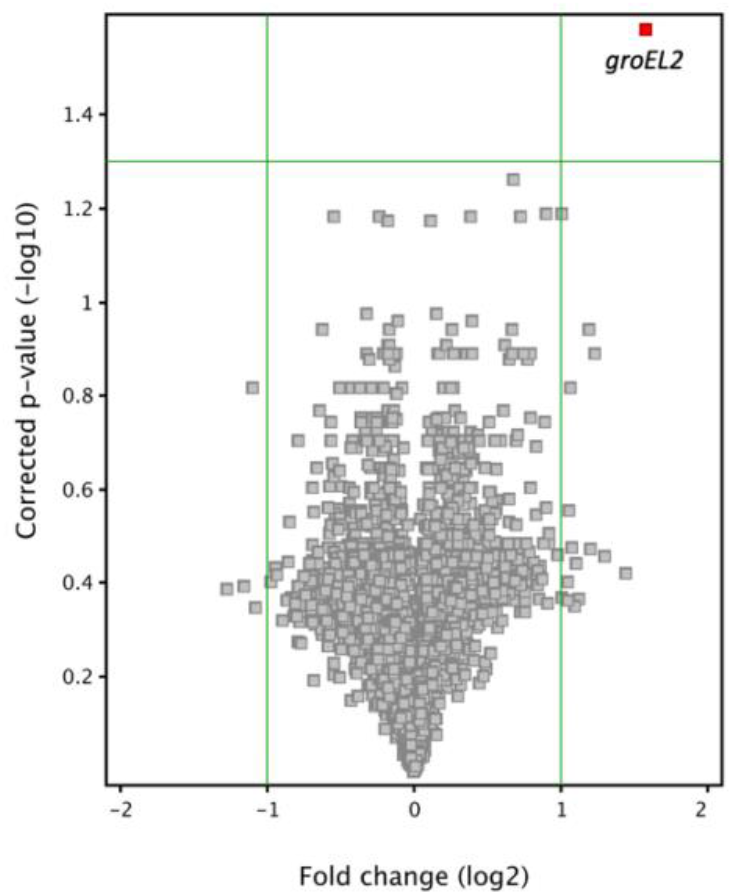
Volcano plot of Δ*sfdS* versus H37Rv upon starvation in PBS. The plot shows *groEL2* as the only gene whose expression level was significantly changed (≥ 2-fold differentially expressed) between Δ*sfdS* and H37Rv.

Strikingly, only a single gene, *groEL2/rv0440*, was significantly differentially expressed, with a 3-fold upregulation in the Δ*sfdS* strain compared to wildtype H37Rv (Fig. 5). To ensure that expression of *groEL2* was restored to wildtype levels in the complemented strain, we performed qRT-PCR on RNA extracted from three biological replicates of wildtype H37Rv, Δ*sfdS* and the complemented strain.

The results confirmed that *groEL2* was significantly upregulated in Δ*sfdS* upon starvation, and that the phenotype could be complemented by providing a copy of the *sfdS* gene in *trans* (Fig. 6). Transcription of *groEL2* is controlled by the HrcA heat shock repressor, which also regulates the *groES-groEL1* and *rv0991c-rv0990c* operons as well as its own expression, i.e. the *hrcA-dnaJ2* operon (28, 29). To determine if the observed changes in *groEL2* expression could be associated with transcriptional control by HrcA, we also measured the expression of the first gene in each of these operons i.e. *groES, rv0991c* and *hrcA* in the three *M. tuberculosis* strains. Similar to that of *groEL2*, the expression of *groES* was significantly upregulated and complemented; *rv0991c* and *hrcA* displayed similar trends, but the changes were not statistically significant (Fig. 6). The role of HrcA in the observed changes therefore remains inconclusive.

**Figure 5:**
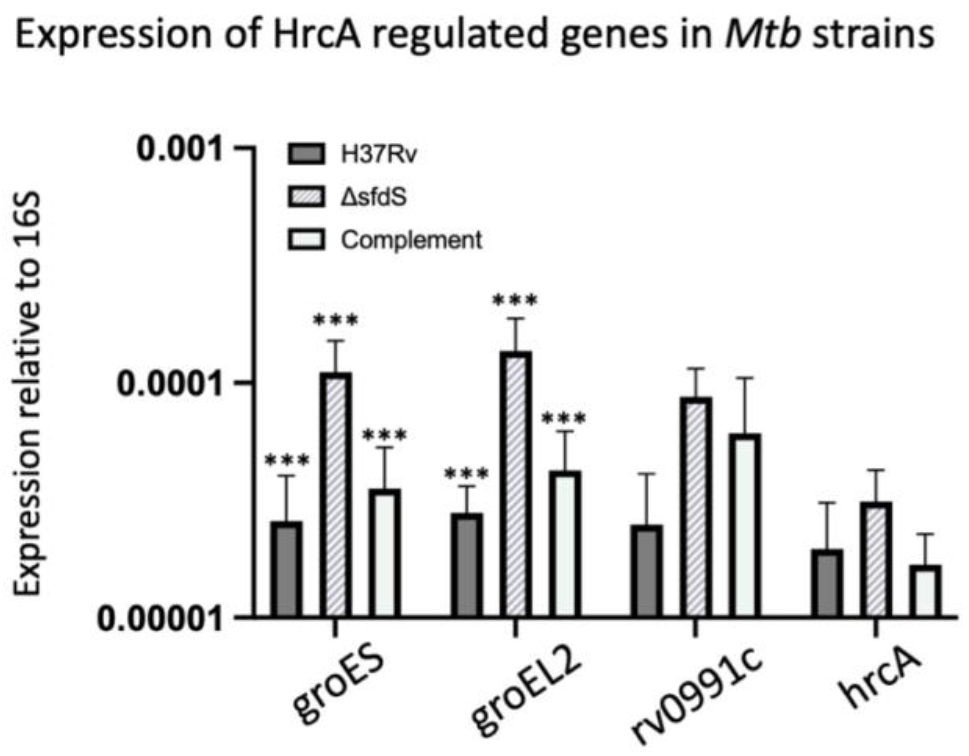
Expression of HrcA-regulated genes in *Mtb* strains. qRT-PCR was performed on total RNA from *M. tuberculosis* H37Rv (wildtype), Δ*sfdS* mutant and its complement. Each bar shows the expression level of the indicated genes normalised to 16S. All data represents the mean and standard deviation of three biological replicates for each strain. *** = p value <0.05 with significance tested using one-way ANOVA.

## DISCUSSION

The *M. tuberculosis* SigF-controlled sRNA SfdS (F6) is well-expressed in exponential phase, but up-regulated by a variety of stresses (6).

In this study we have shown that SfdS is also highly upregulated after three weeks of murine infection, and to an even higher extent in the PBS starvation model at 24 and 96 hours. This suggests that SfdS plays a role in the early stages of pathogenesis but possibly a more significant one for survival in nutrient deficient environments such as those encountered in human granulomas in later stages of infection. *M. tuberculosis* may resuscitate after periods of low metabolic activity (27, 30, 31), and using the Wayne model of non-replicating persistence, we found that recovery of Δ*sfdS* from NRP-2 was impaired compared to wildtype and complemented strains, although initial survival was unchanged. The fact that the complemented strain displayed an intermediate phenotype, may be due to *cis*-regulatory effects associated with the deletion, the different genomic location of the complementing SfdS or the point mutations identified with the WGS.

Our results indicate that starvation for 24 hours leads to the upregulation of HrcA-controlled chaperonins *groES and groEL2*, in the Δ*sfdS* strain compared to the wildtype and complemented *M. tuberculosis* strains, while the expression of the remaining HrcA-regulated operons (*rv0991c-0990c* and *hrcA-dnaJ2*) were not significantly changed. However, regulation of HrcA expression and activity is highly complex and the lack of a significant change of *hrcA* mRNA levels in the Δ*sfdS* strain does not exclude a role for this regulator in the observed response.

Prediction of potential mRNA targets of SfdS using the TargetRNA2 webserver (25) with default settings against the H37Rv genome resulted in 27 potential targets. However, none of these could be associated with the partial heat shock signature that we observed, and we conclude that the observed changes likely represent downstream effects from other, as yet unidentified targets.

Nevertheless, the fact that SfdS appears to repress chaperonin expression does offer a potential explanation for the previously observed slow growth associated with SfdS overexpression (6), namely an untimely repression of these essential chaperonins, which could lead to reduced protein synthesis. This also suggests that the overexpression strain may be hyper-sensitive to heat shock.

RNA is a ubiquitous temperature-sensing molecule, and several heat shock response mechanisms employ RNA, including the eukaryotic heat shock transcription factor 1 (HSF1), *Eschericia coli rpoH* and *Deinococcus radiodurans* sRNA DnrH, the latter which activates the expression of Hsp20 (32–34). Further analysis using pulsed rather than constitutive overexpression or deletion may provide more clues on the link between SfdS and heat shock.

Infection of BMDMs for 7 days with H37Rv, Δ*sfds* and complement revealed no attenuation of the deletion strain. Similarly, we observed no difference in the CFUs recovered from the lungs and spleens of BALB/c mice after 143 days of infection. As SfdS expression is not only directed by SigF, but also dominant in terms of promoter occupancy, one might expect some parallels between infection with Δ*sfdS* and Δ*sigF*. Indeed, a lack of attenuation has been observed in a study of a Δ*sigF* strain in human monocyte derived macrophages, while late stages of infection and time-to-death experiments result in attenuation of a CDC1551 Δ*sigF* strain (18, 35). Given that starvation is a potent inducer of SigF/SfdS expression, it seems possible that a more long-term infection and/or the ability to form granulomas, are required to observe strong phenotypes associated with the deletion of *sfdS*. Moreover, the mouse model of infection does not replicate all aspects of an *M. tuberculosis* infection in humans, one key difference being the lack of granuloma formation in a standard mouse model such as BALB/c (36). Granulomas provide a unique niche for *M. tuberculosis* in which oxygen and nutrients are limited, the latter being the strongest inducer of SfdS expression in our study. A different animal model e.g. using C3HeB/FeJ mice, which develop necrotic lung granulomas, may provide more answers (37). Our results point towards a role for SfdS during periods of low metabolic activity similar to those of cold shock and may be associated with nutrient starvation conditions such as those found in human granulomas in later stages of infection, and it remains a possibility that SfdS plays a role in surviving and resuscitating from this environment.

## ACCESSION NUMBERS

Array design is available in BμG@Sbase (Accession No. A-BUGS-41; http://bugs.sgul.ac.uk/A-BUGS-41) and ArrayExpress (Accession No. A-BUGS-41). Microarray data have been deposited in ArrayExpress (Accession number E-MTAB-9327).

## ACKNOWLEDGEMENTS

We thank Ruben Hartkoorn and Stewart Cole for the H37Rv Δ*sigF* mutant, the Biological Service staff at NIMR, Mill Hill for help with maintaining mice, the BμG@S (the Bacterial Microarray Group at St. George’s, University of London) for supplying *M. tuberculosis* microarrays, Finn Werner for critical reading of the manuscript and Douglas Young for support and mentoring.

## REFERENCES

1. Arnvig KB, Comas I, Thomson NR, Houghton J, Boshoff HI, Croucher NJ, Rose G, Perkins TT, Parkhill J, Dougan G, Young DB. 2011. Sequence-based analysis uncovers an abundance of non-coding RNA in the total transcriptome of Mycobacterium tuberculosis. PLoS Pathog 7:e1002342.

2. Cortes T, Schubert OT, Rose G, Arnvig KB, Comas I, Aebersold R, Young DB. 2013. Genome-wide mapping of transcriptional start sites defines an extensive leaderless transcriptome in Mycobacterium tuberculosis. Cell Rep 5:1121–31.

3. Houghton J, Cortes T, Schubert O, Rose G, Rodgers A, De Ste Croix M, Aebersold R, Young DB, Arnvig KB. 2013. A small RNA encoded in the Rv2660c locus of Mycobacterium tuberculosis is induced during starvation and infection. PLoS One 8:e80047.

4. Miotto P, Forti F, Ambrosi A, Pellin D, Veiga DF, Balazsi G, Gennaro ML, Di Serio C, Ghisotti D, Cirillo DM. 2012. Genome-wide discovery of small RNAs in Mycobacterium tuberculosis. PLoS One 7:e51950.

5. Shell SS, Wang J, Lapierre P, Mir M, Chase MR, Pyle MM, Gawande R, Ahmad R, Sarracino DA, Ioerger TR, Fortune SM, Derbyshire KM, Wade JT, Gray TA. 2015. Leaderless Transcripts and Small Proteins Are Common Features of the Mycobacterial Translational Landscape. PLoS Genet 11:e1005641.

6. Arnvig KB, Young DB. 2009. Identification of small RNAs in Mycobacterium tuberculosis. Mol Microbiol 73:397–408.

7. DiChiara JM, Contreras-Martinez LM, Livny J, Smith D, McDonough KA, Belfort M. 2010. Multiple small RNAs identified in Mycobacterium bovis BCG are also expressed in Mycobacterium tuberculosis and Mycobacterium smegmatis. Nucleic Acids Res 38:4067–78.

8. Gerrick ER, Barbier T, Chase MR, Xu R, Francois J, Lin VH, Szucs MJ, Rock JM, Ahmad R, Tjaden B, Livny J, Fortune SM. 2018. Small RNA profiling in Mycobacterium tuberculosis identifies MrsI as necessary for an anticipatory iron sparing response. Proc Natl Acad Sci U S A 115:6464–6469.

9. Mai J, Rao C, Watt J, Sun X, Lin C, Zhang L, Liu J. 2019. Mycobacterium tuberculosis 6C sRNA binds multiple mRNA targets via C-rich loops independent of RNA chaperones. Nucleic Acids Res 47:4292–4307.

10. Moores A, Riesco AB, Schwenk S, Arnvig KB. 2017. Expression, maturation and turnover of DrrS, an unusually stable, DosR regulated small RNA in Mycobacterium tuberculosis. PLoS One 12:e0174079.

11. Pelly S, Bishai WR, Lamichhane G. 2012. A screen for non-coding RNA in Mycobacterium tuberculosis reveals a cAMP-responsive RNA that is expressed during infection. Gene 500:85–92.

12. Salina EG, Grigorov A, Skvortsova Y, Majorov K, Bychenko O, Ostrik A, Logunova N, Ignatov D, Kaprelyants A, Apt A, Azhikina T. 2019. MTS1338, A Small Mycobacterium tuberculosis RNA, Regulates Transcriptional Shifts Consistent With Bacterial Adaptation for Entering Into Dormancy and Survival Within Host Macrophages. Front Cell Infect Microbiol 9:405.

13. Solans L, Gonzalo-Asensio J, Sala C, Benjak A, Uplekar S, Rougemont J, Guilhot C, Malaga W, Martin C, Cole ST. 2014. The PhoP-dependent ncRNA Mcr7 modulates the TAT secretion system in Mycobacterium tuberculosis. PLoS Pathog 10:e1004183.

14. Lovewell RR, Sassetti CM, VanderVen BC. 2016. Chewing the fat: lipid metabolism and homeostasis during M. tuberculosis infection. Curr Opin Microbiol 29:30–6.

15. Hartkoorn RC, Sala C, Uplekar S, Busso P, Rougemont J, Cole ST. 2012. Genome-wide definition of the SigF regulon in Mycobacterium tuberculosis. J Bacteriol 194:2001–9.

16. Betts JC, Lukey PT, Robb LC, McAdam RA, Duncan K. 2002. Evaluation of a nutrient starvation model of Mycobacterium tuberculosis persistence by gene and protein expression profiling. Mol Microbiol 43:717–31.

17. DeMaio J, Zhang Y, Ko C, Young DB, Bishai WR. 1996. A stationary-phase stress-response sigma factor from Mycobacterium tuberculosis. Proc Natl Acad Sci U S A 93:2790–4.

18. Geiman DE, Kaushal D, Ko C, Tyagi S, Manabe YC, Schroeder BG, Fleischmann RD, Morrison NE, Converse PJ, Chen P, Bishai WR. 2004. Attenuation of late-stage disease in mice infected by the Mycobacterium tuberculosis mutant lacking the SigF alternate sigma factor and identification of SigF-dependent genes by microarray analysis. Infect Immun 72:1733–45.

19. Michele TM, Ko C, Bishai WR. 1999. Exposure to antibiotics induces expression of the Mycobacterium tuberculosis sigF gene: implications for chemotherapy against mycobacterial persistors. Antimicrob Agents Chemother 43:218–25.

20. Hinds J, Mahenthiralingam E, Kempsell KE, Duncan K, Stokes RW, Parish T, Stoker NG. 1999. Enhanced gene replacement in mycobacteria. Microbiology (Reading) 145 (Pt 3):519–527.

21. Gopaul KK, Brooks PC, Prost JF, Davis EO. 2003. Characterization of the two Mycobacterium tuberculosis recA promoters. J Bacteriol 185:6005–15.

22. Springer B, Master S, Sander P, Zahrt T, McFalone M, Song J, Papavinasasundaram KG, Colston MJ, Boettger E, Deretic V. 2001. Silencing of oxidative stress response in Mycobacterium tuberculosis: expression patterns of ahpC in virulent and avirulent strains and effect of ahpC inactivation. Infect Immun 69:5967–73.

23. Li H, Durbin R. 2009. Fast and accurate short read alignment with Burrows-Wheeler transform. Bioinformatics 25:1754–60.

24. Li H, Handsaker B, Wysoker A, Fennell T, Ruan J, Homer N, Marth G, Abecasis G, Durbin R, Genome Project Data Processing S. 2009. The Sequence Alignment/Map format and SAMtools. Bioinformatics 25:2078–9.

25. Kery MB, Feldman M, Livny J, Tjaden B. 2014. TargetRNA2: identifying targets of small regulatory RNAs in bacteria. Nucleic Acids Res 42:W124–9.

26. Ehlers S, Schaible UE. 2012. The granuloma in tuberculosis: dynamics of a host-pathogen collusion. Front Immunol 3:411.

27. Wayne LG, Hayes LG. 1996. An in vitro model for sequential study of shiftdown of Mycobacterium tuberculosis through two stages of nonreplicating persistence. Infect Immun 64:2062–9.

28. Hakiem OR, Parijat P, Tripathi P, Batra JK. 2020. Mechanism of HrcA function in heat shock regulation in Mycobacterium tuberculosis. Biochimie 168:285–296.

29. Stewart GR, Wernisch L, Stabler R, Mangan JA, Hinds J, Laing KG, Young DB, Butcher PD. 2002. Dissection of the heat-shock response in Mycobacterium tuberculosis using mutants and microarrays. Microbiology (Reading) 148:3129–3138.

30. Rosser A, Stover C, Pareek M, Mukamolova GV. 2017. Resuscitation-promoting factors are important determinants of the pathophysiology in Mycobacterium tuberculosis infection. Crit Rev Microbiol 43:621–630.

31. Salina EG, Waddell SJ, Hoffmann N, Rosenkrands I, Butcher PD, Kaprelyants AS. 2014. Potassium availability triggers Mycobacterium tuberculosis transition to, and resuscitation from, non-culturable (dormant) states. Open Biol 4.

32. Shamovsky I, Ivannikov M, Kandel ES, Gershon D, Nudler E. 2006. RNA-mediated response to heat shock in mammalian cells. Nature 440:556–60.

33. Xue D, Chen Y, Li J, Han J, Liu Y, Jiang S, Zhou Z, Zhang W, Chen M, Lin M, Ongena M, Wang J. 2019. Targeting Hsp20 Using the Novel Small Non-coding RNA DnrH Regulates Heat Tolerance in Deinococcus radiodurans. Front Microbiol 10:2354.

34. Yura T. 2019. Regulation of the heat shock response in Escherichia coli: history and perspectives. Genes Genet Syst 94:103–108.

35. Chen P, Ruiz RE, Li Q, Silver RF, Bishai WR. 2000. Construction and characterization of a Mycobacterium tuberculosis mutant lacking the alternate sigma factor gene, sigF. Infect Immun 68:5575–80.

36. Orme IM, Basaraba RJ. 2014. The formation of the granuloma in tuberculosis infection. Semin Immunol 26:601–9.

37. Harper J, Skerry C, Davis SL, Tasneen R, Weir M, Kramnik I, Bishai WR, Pomper MG, Nuermberger EL, Jain SK. 2012. Mouse model of necrotic tuberculosis granulomas develops hypoxic lesions. J Infect Dis 205:595–602.

