## Supplementary data for "The *Mycobacterium tuberculosis* sRNA F6 modifies expression of essential chaperonins, GroEL2 and GroES"

### SUPPLEMENTARY MATERIAL

#### Table S1: Plasmids used in this study

| Plasmid | Relevant Characteristic | Source or Reference |
|---|---|---|
| pBackbone | Mycobacterial suicide vector (kanamycin$^R$ and ampicillin$^R$) | (1) |
| pKP186 | Integrating mycobacterial cloning vector that does not contain intergrase (kanamycin$^R$) | (2) |
| PBSInt | Mycobacterial suicide vector containing integrase, electroporated in conjunction with pKP186 and its derivatives. | (3) |
| pJHP04 | Targeting plasmid for removal of F6 in *M. tuberculosis*. pBackbone containing F6 5' and 3' flanking regions and the sacB/lacZ cassette | This Study |
| pJHP06 | F6 complementing plasmid. pKP186 derivative containing 448bp coordinates 293428-293876 | This Study |

#### Table S2: Oligos used in this study

| Name | Sequence (5'-3') | Description |
|---|---|---|
| F6RevXbaI | GGTCTAGACGAGTGATCGGG | F6 targeting plasmid |
| F6ForXbaI | GGTCTAGATGGGCTTGCCC | F6 targeting plasmid |
| F6RevSDM | GGGGCAAGCCCAAAAAGATCTAGACCGAGTGATC GGGTACCC | F6 targeting plasmid |
| F6ForSDM | GGGTACCCGATCACTCGGTCTAGATCTTTTTGGGC TTGCCCC | F6 targeting plasmid |
| F6compF | GAAAAGCTTGCCGCTGTTGACCAG | F6 complement plasmid |
| F6compR | CGGATCCCTGCGCGGGCTGA | F6 complement plasmid |
| F6nrt | CGGATAGCCCCGTGTTGTTGTCTGACCTGTCTC | F6 Northern probe template |
| 16sTqmF | TCCCGGGCCTTGTACACA | qRT-PCR |
| 16sTqmR | CCACTGGCTTCGGGTGTTA | qRT-PCR |
| F6TqmF | GGATAGCCCCGTGTTGTTG | qRT-PCR |
| F6TqmR | GGGATTGCCCCGCATT | qRT-PCR |
| Rv0440TqmF | CGTCGTCCTGGAAAAGAAGTG | qRT-PCR |
| Rv0440TqmR | GGTCTTCTTGGCTACCTCTTTGAC | qRT-PCR |

| | | |
|---|---|---|
| Rv3418cTqmF | CGTTGCGGAGGGTGACA | qRT-PCR |
| Rv3418cTqmR | TCCTCGCCGTTGTACTTGATC | qRT-PCR |
| Rv0990cTqmF | GGCCGCGCACGATCT | qRT-PCR |
| Rv0990cTqmR | CGTTTTTCCAGCCTGACATCA | qRT-PCR |
| Rv0991cTqmF | TTCAAAGGCACCGGCTTCTA | qRT-PCR |
| Rv0991cTqmR | TGGTCTGGCTCTTGGACTTCTT | qRT-PCR |

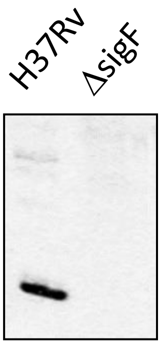


**Fig. S1: No F6 expression in Δ*sigF* mutant**. 10 μg each of H37Rv and Δ*sigF* total RNA was separated on a 10% denaturing acrylamide gel and transferred onto Brightstar-Plus nylon membrane (Ambion) by electroblotting. RNA was UV cross-linked to the membrane and stained with 0.3 M sodium acetate/0.03% methylene blue to verify transfer. 32P-labelled riboprobes were synthesised using the mirVana Probe construction Kit (Ambion) and 32P-UTP (800 mCi/mmol, PerkinElmer) with the template oligo listed Table S2, and hybridised to the membranes overnight in UltraHyb (Invitrogen).

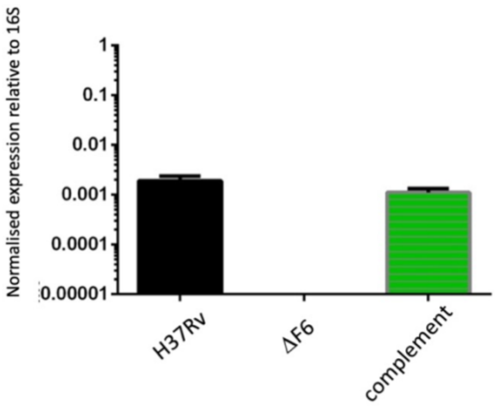


**Fig. S2 Expression of F6 in different strains**

RNA from 7H9 cultures of H37Rv, ΔF6 and the complemented strain was harvested in exponential phase and F6 levels measured by qRT-PCR normalised to 16S rRNA. Data represents the average and standard deviation of three biological replicates for each strain.

### References for supplementary data


1. Gopaul KK, Brooks PC, Prost JF, Davis EO. 2003. Characterization of the two Mycobacterium tuberculosis recA promoters. J Bacteriol 185:6005-15.
2. Rickman L, Scott C, Hunt DM, Hutchinson T, Menendez MC, Whalan R, Hinds J, Colston MJ, Green J, Buxton RS. 2005. A member of the cAMP receptor protein family of transcription regulators in Mycobacterium tuberculosis is required for virulence in mice and controls transcription of the rpfA gene coding for a resuscitation promoting factor. Mol Microbiol 56:1274-86.
3. Springer B, Master S, Sander P, Zahrt T, McFalone M, Song J, Papavinasasundaram KG, Colston MJ, Boettger E, Deretic V. 2001. Silencing of oxidative stress response in Mycobacterium tuberculosis: expression patterns of ahpC in virulent and avirulent strains and effect of ahpC inactivation. Infect Immun 69:5967-73.
